# Flu Virus Infection in Juvenile Mice Leads to Lifelong Multi-Organ Damage and Parkinsonian Pathological Changes in Aging

**DOI:** 10.64898/2025.12.24.696086

**Authors:** Ruina You, Cuicui Liu, Xun Liu, Xinglian Wang, Zhuoya Xu, Qian Huang, Guolin Dong, Chaoqun Liang, Na Wang, Zhichao Miao, Taijiao Jiang, Dayan Wang, Lei Deng, Yousong Peng

**Affiliations:** Bioinformatics Center, College of Biology, Hunan Provincial Key Laboratory of Medical Virology, Hunan University, Changsha 410082, China; Anhui Medical University, Hefei, China, Department of Otolaryngology Head and Neck Surgery, The Second Affiliated Hospital of Anhui Medical University, Hefei, China; Bengbu Medical University, Bengbu, China, Obstetrics and Gynecology, The Second People’s Hospital of Wuhu, Wuhu, China; Guangzhou National Laboratory, Guangzhou 510005, China; National Institute for Viral Disease Control and Prevention, China CDC, Beijing, 102206, China

## Abstract

The widespread prevalence of Long COVID underscores the potential for viral infections to exert prolonged effects on hosts long after initial recovery. Despite this recognition, the long-term and even lifelong consequences of viral infections remain poorly understood. In this study, we infected juvenile mice with influenza viruses and systematically examined the lifelong effects (from 4 to 450 days post-infection) of these infections. Pathological analysis revealed persistent lifelong damage in multiple organs, including chronic lung inflammation and fibrosis, along with significant cardiac and renal pathology. Viral infection led to substantial neuronal loss in key brain regions of mice (hippocampal CA1, CA3, and striatum) during middle and old age. Strikingly, we identified two hallmark pathological features of Parkinson’s disease - dopaminergic neuron degeneration and Lewy body–like α-synuclein inclusions - in middle and aged infected mice. Transcriptomic profiling demonstrated sustained upregulation of inflammation-related genes in both lung and blood tissues, correlating with observed pathological changes. Lung-derived secreted proteins, encoded by differentially expressed genes, may mediate cross-organ communication affecting cardiac, renal, and neurological function. Brain transcriptome analysis at three months post-infection revealed downregulation of neurodevelopmental genes, potentially contributing to subsequent neuronal loss and Parkinsonian pathology. These findings collectively suggest that pulmonary influenza infection can induce systemic multi-organ effects through both secretory pathways and chronic inflammatory responses. The study offers crucial insights for developing more effective strategies to prevent and manage infectious diseases and their long-term sequelae.

## Main

Viruses are one of the most common pathogens to humans and can cause acute, chronic, or latent infections in diverse tissues. Acute viral infections, such as influenza and COVID-19, typically cause rapid-onset illness^1^. In contrast, chronic viral infections persist within the host, often establishing lifelong latency or sustained replication while evading immune clearance. These infections contribute to ongoing inflammation, tissue damage, and cancer development^2^. Notable examples include hepatitis B virus (HBV), human immunodeficiency virus (HIV), and human papillomavirus (HPV), all of which impose substantial long-term disease burdens^3–5^. Unlike the chronic or latent infections, recent burst of Long COVID that is characterized by symptoms such as fatigue, cognitive dysfunction, and respiratory issues that persist for months after the initial infection, has brought to light the potential for viral infections to have long-lasting effects on human health after recovery from the acute phase of illness.

While the acute phase of virus infection has been extensively studied, the long-term consequences of such infections remain poorly understood. A recent study by Levine et al. showed that exposures to viruses such as influenza virus and EBV were associated with an increased risk of neurodegeneration up to 15 years after infection by analysis of cohort data in FinnGen and UK Biobank^6^. Conducting long-term studies in human populations is particularly challenging due to the multitude of confounding factors that can influence outcomes, including pre-existing health conditions, lifestyle factors, and environmental exposures. In contrast, animal models have provided valuable insights into the long-term effects of virus infections. For instance, studies have shown that influenza A virus (IAV) infection in mice can lead to long-term neuroinflammation, with effects on hippocampal neuron morphology and function persisting for up to three months post-infections^7^. This long-term neuroinflammation was associated with impairments in spatial memory formation and changes in dendritic spine density in the hippocampus. Additionally, the infection led to increased activation of microglia, the brain’s resident immune cells, and altered synaptic plasticity. These findings suggest that the effects of influenza infection can extend well beyond the acute phase and may have significant implications for cognitive function and neuronal health^8,9^. However, the effects of influenza virus infection over even longer periods, including lifelong impacts, remain largely unexplored.

Viral infections can impact not only the directly infected tissues but also multiple organs and systems throughout the body^10^. For instance, SARS-CoV-2, the virus responsible for COVID-19, has been shown to cause symptoms across various systems, including the cardiovascular, respiratory, and nervous systems. The virus can directly breach common sites of infection, such as the respiratory tract, to cause systemic infection. A recent study by Alon et al. showed that the influenza virus can cause damage to multiple organs by entering into bloodstream following lung infections^11^. Besides, the virus can also trigger systemic inflammatory responses through the immune system, which can affect the function of other organs. This systemic inflammation can lead to a cascade of effects, including the release of pro-inflammatory cytokines and chemokines that can influence distant tissues. Particularly concerning is the impact on the central nervous system, where viral-induced immune activation can lead to neuroinflammation. This neuroinflammation can disrupt the blood-brain barrier, allowing immune cells and inflammatory molecules to enter the brain, potentially contributing to long-term cognitive impairments and neurodegenerative diseases^12,13^. Despite this advancement, the molecular mechanism through which virus cause multiple-organ damage remains unclear.

This study aims to explore the life-long effects of influenza virus infection using a mouse model. The juvenile mice (6~8 weeks) were infected with the mouse-adapted HK68 strain (Figure 1a). Then, the mouse tissues including lung, blood, brain, heart and kidney were collected on 4, 20, 90, 270 and 450 Dpi for both virus infection group and PBS group. The tissue samples were divided into two parts for processing: one part was used for transcriptomic analysis based on RNA-sequencing, and the other part was used for histological and pathological analyses which includes Hematoxylin and eosin (H&E) staining, Nissl staining, Masson’s trichrome staining, and immunofluorescence double-labeling (Tyrosine Hydroxylase (TH) and α-synuclein).

**Figure 1.**
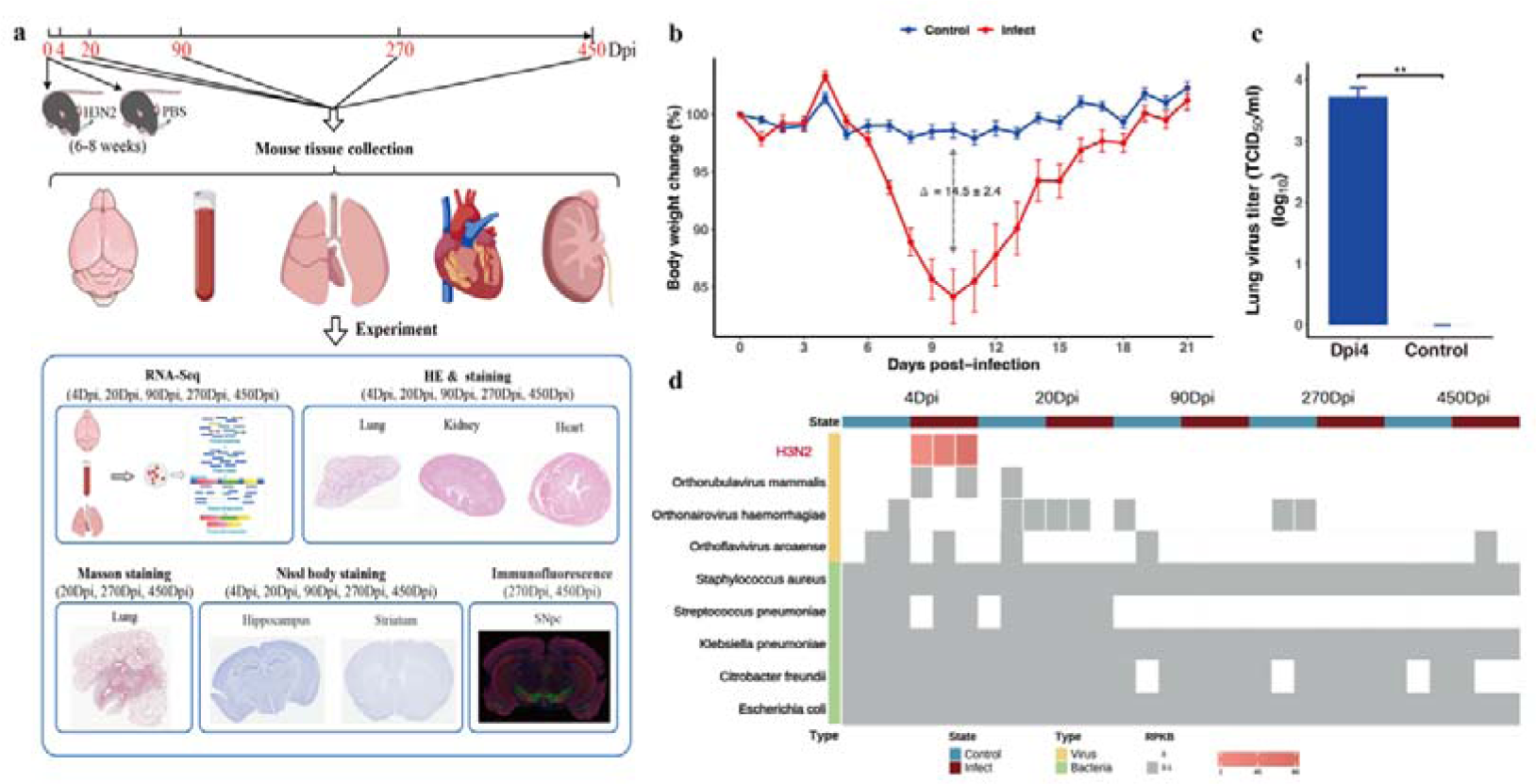
Overview of the study and the detection of pathogens in mice. (**a**) Overview of the study. (**b**) Body weight changes of mice in viral infection (red) and PBS group (blue) from 0 to 21Dpi. The curves showed the average and standard errors of daily body weight of mice. The delta showed the largest difference between the viral infection and control groups. (**c**) Detection of influenza virus load in mouse lung at 4 Dpi. (**d**) Detection of viruses and bacteria in mouse lung at different time points after infections based on RNA-Seq data. The pathogen abundance was indicated based on the legend at the bottom. The gray refers to abundance of smaller than 0.1 RPKB (see Methods).

### Validation of influenza virus infection in mouse lung

The body weight of mice in viral infection group decreased much from 5~10 Dpi and was lowest at 10 Dpi with an average decrease of 14.5% compared to the PBS group (Figure 1b). The mice gradually regained body weight, reaching near-normal levels around 20 Dpi. Analysis of the viral load by TCID_50_ showed that the mouse lung in the infection group had an average of 3.8 log_10_ TCID_50_/ml at 4 Dpi, while the PBS group had no virus detected at this time (Figure 1c). We further analyzed the RNA-Seq data to identify possible pathogens in mice. As shown in Figure 1d and Table S1, a high abundance of influenza virus was identified in mouse lung of virus-infection group at 4 Dpi, with an average of 81.5 RPKB (see Methods). They were not detected in lung at other time points (20 Dpi, 90 Dpi, 270 Dpi and 450 Dpi), and were also not detected in blood and brain at any time point. In the PBS group, no influenza virus was detected in any tissue (lung, blood or brain) at five time points. Besides the influenza virus, there were three viruses detected in mice with extremely low abundance (<0.01 PRKB), including Orthorubulavirus mammalis, Orthonairovirus haemorrhagiae, and Orthoflavivirus aroaense (Figure 1d and Table S1). Several bacterial pathogens were also detected in nearly all samples of both viral infection and PBS groups with extremely low abundance (< 0.1 RPKB in most cases) (Figure 1d and Table S1), such as Staphylococcus aureus, Streptococcus pneumoniae and Klebsiella pneumoniae. They were more likely to be contaminated since they were commonly detected in the sequencing. These results suggest that the influenza virus was the only dominant pathogen in viral infection group.

### Influenza virus infection causes life-long damage and pulmonary fibrosis in mouse lungs

Then, we analyzed the pathological changes of mice lung after influenza virus infection by H&E staining. The results showed that the mice lung exhibited typical features of exudative inflammation at 4 Dpi (see red arrows), characterized by thickened alveolar septa, diffuse inflammatory cell infiltration in the interstitium, and proteinaceous exudative edema in the alveolar spaces (Figure 2a). The inflammation continued to exist even when the mice had restored from infection, although the extent had decreased. No obvious injury was observed in the PBS group except at 450 Dpi, when a low level of injury was observed, such as increased inflammatory cell infiltration and alveolar thickening. We then quantified the extent of injury in mice lung from six aspects including alveolar edema, the number of cell fragments, inflammatory cell infiltration, infiltration of alveolar cells, area of alveolar hemorrhage and pulmonary alveolar thickening (Table S2). A score with 0~4 was assigned to each aspect and the sum of these scores was used to reflect the overall extent of injury in lung (see Methods). As shown in Figure 2e, the overall lung injury score was highest in viral infection mice lung at 4 Dpi (total average score =21.00±1.95). Specifically, the average score for alveolar edema, the number of cell fragments, inflammatory cell infiltration, infiltration of alveolar cells, area of alveolar hemorrhage, and pulmonary alveolar thickening was 3.50, 3.75, 4.12, 3.88, 3.50 and 2.25, respectively (Table S2). This suggested serious inflammation in the viral infection group at 4 Dpi. In contrast, a very low lung injury score (1.38±0.38) was observed in the PBS group at the same time. After recovered from viral infection, the lung injury score in the lung decreased much at 20 Dpi. Interestingly, even when a long time from recovery (90 Dpi), there were still a median level of inflammation with the lung injury score of 9.62. The lung injury score in viral infection mice continued to decrease from 90 Dpi to 270 Dpi and 450 Dpi when a low level of injury was observed. In contrast, the PBS group showed increased injury from 90 Dpi to 270 Dpi, reaching levels comparable to those in the viral infection group by 450 Dpi, at which point both groups exhibited inflammatory cell infiltration and alveolar thickening (Table S2).

**Figure 2.**
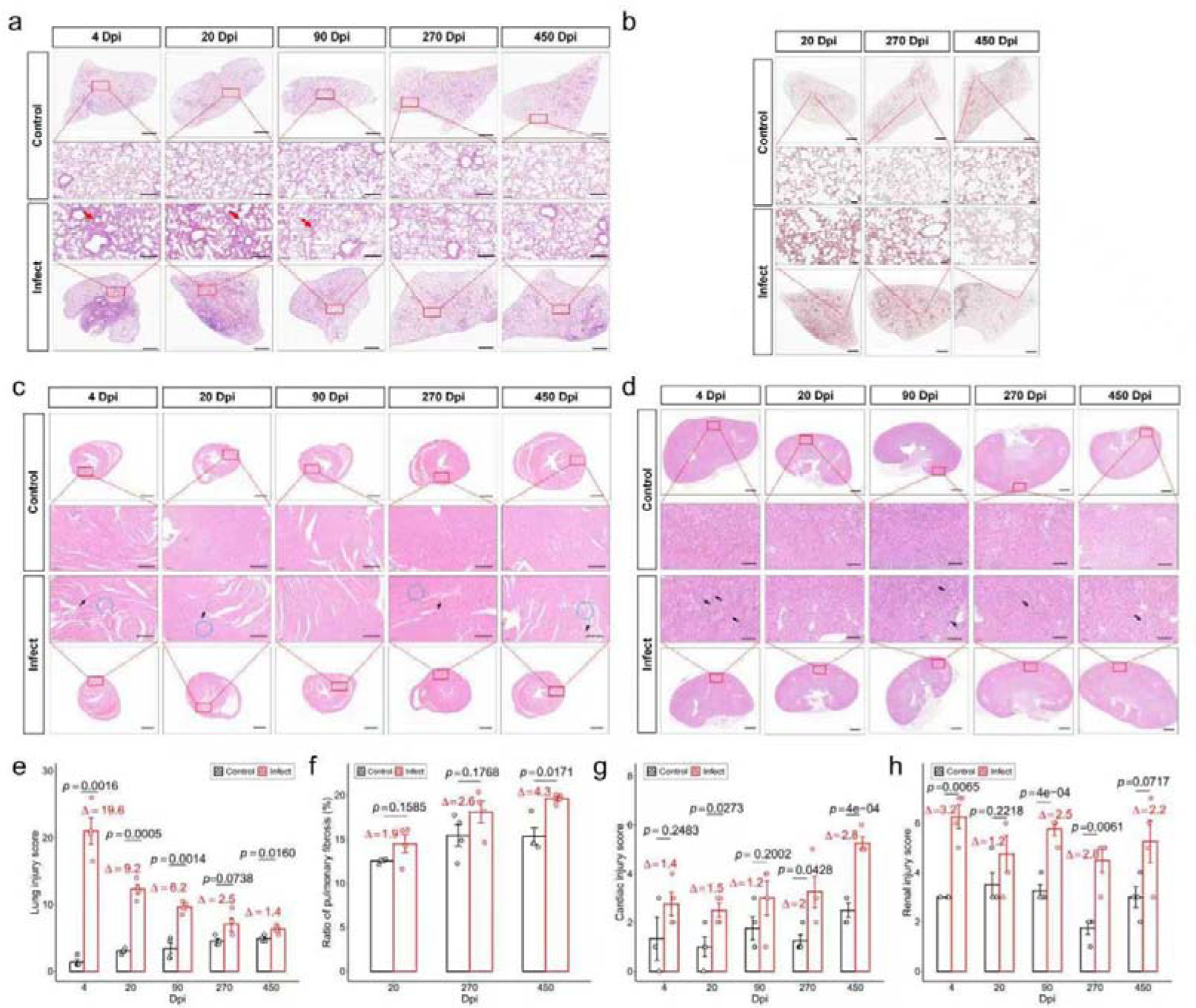
Pathological changes in the lung, heart, and kidneys of mice following influenza virus infection, assessed by H&E staining (lung, heart, and kidneys) and Masson staining (lung only). (**a**) and (**b**) refer to the lung images at different time points by H&E staining (20×, Scale bar = 1 mm for low magnification; 200×, Scale bar = 200 μm for high magnification) and Masson staining (20×, Scale bar = 1 mm for low magnification; 250×, Scale bar = 50 μm for high magnification), respectively. (**c**) and (**d**) refer to the images of the heart and kidney, respectively, at five time points by H&E staining (10×, Scale bar = 1 mm for low magnification; 100×, Scale bar = 100 μm for high magnification). (**e**) and (**f**) refer to the lung injury score and the alveolar collagen deposition area ratio, respectively, at different time points. (**g**) and (**h**) refer to the pathology scores of the heart and kidney, respectively. The bars show the average and the standard errors, and the delta indicates the average difference between the infection group and the PBS group. n = 4 mice per group. Representative images are shown.

We then perform Masson’s trichrome staining to investigate the mice pulmonary fibrosis in mice following viral infections. As shown in Figure 2b, markedly increased collagen deposition (see the blue area) was observed in the alveoli of viral infection mice compared to those of PBS group at 270 Dpi and 450 Dpi. Quantitative analysis showed that the alveolar collagen deposition area ratio in both virus-infected group and PBS group were smaller than 15% at 20 Dpi (Figure 2f and Table S3), suggesting no fibrosis in the mice lung; the ratios in both groups increased to be greater than 15% at 270 Dpi, suggesting the emergence of pulmonary fibrosis in the adult mice; as the mice grown into the old, the ratio in the PBS group keep stable, while that in the virus-infected group continued to increase, indicating a progressive pulmonary fibrosis. The alveolar collagen deposition area ratio in the virus-infected group was consistently higher than that in the PBS group, although statistical difference was only observed at 450 Dpi (p=0.0171) (Figure 2f).

### Influenza virus infections cause long-term damage to the heart and kidneys

Next, we examined the pathological damage to the heart and kidneys induced by influenza virus infection using H&E staining. As shown in Figure 2c, significant pathological damage was observed in the mice heart at 4 Dpi, including changes in cardiac inflammatory response and myocardial degeneration. This pathological damage persisted and became more pronounced at 450 Dpi (black arrows and blue dashed circles). In contrast, mice in the PBS group exhibited only minor pathological damage in their hearts during old age (450 Dpi). Next, we quantitatively assessed the pathological damage in mouse hearts from three aspects: inflammatory response, myocardial degeneration, and fibrosis (Table S4). Each aspect was scored on a scale of 0 to 4, and the sum of the three scores represented the total cardiac pathology score (see Methods). For any aspect, the infection group generally scored higher than the PBS group in most cases (Table S4). For the total score, at all five time points, the infection group’s scores were significantly higher than those of the PBS group (Figure 2g), though statistically significant differences were only observed at 20, 270, and 450 Dpi (p-values: 0.0273, 0.0428, and 4e-04, respectively). Notably, at 270 and 450 Dpi, the infection group’s scores were twice as high as those of the PBS group. These findings suggest that the detrimental effects of influenza virus infection on the heart become significantly exacerbated in middle and old age.

Influenza virus infection caused significant damage to the mice kidney (Figure 2d), including changes in glomerular inflammatory cell infiltration and the proportion of sclerotic glomeruli (indicated by arrows). We quantitatively assessed renal pathological damage from two aspects: concurrent glomerulosclerosis and inflammatory infiltration (Table S5), each scored on a scale of 0 to 4. The sum of these two scores represented the total renal pathology score (see Methods). Interestingly, influenza virus infection induced a triphasic pattern of kidney injury: acute phase, remission phase, and chronic progression phase (Figure 2h). During active viral infections (4 Dpi), the kidneys exhibited acute inflammation, with pathological scores double those of the control group; as the mice recovered from infection (20 Dpi), renal damage partially resolved, though pathological scores remained elevated compared to controls (without statistical significance); by 90 Dpi and later, renal injury resurged to levels comparable to the acute phase, followed by a gradual decline—yet remaining persistently higher than controls. These results indicate that, although the mice fully recovered from acute influenza infection, the kidneys sustained long-term pathological damage.

### Influenza virus infection leads to neuronal loss in the hippocampus and striatum of mice

Next, we analyzed the impact of influenza virus infection on the central nervous system (CNS) of mice. Nissl staining revealed that on 4 Dpi, the neuron density in the whole CA1 region of the hippocampus in viral infection group was comparable to that of the PBS group, with an average of 106/mm² and 104/mm², respectively (Figure 3a&b). However, from 20 to 450 Dpi, the infected mice exhibited a reduced density of neurons in the hippocampal CA1 region compared to the PBS group, with obvious differences (delta, see the black triangles) observed particularly at 270 Dpi (delta = 19.8) and 450 Dpi (delta = 14.0). For example, the average neuron density in the infected group was 79.8/mm² at 270 Dpi, while that of the PBS group was 99.5/mm². In the CA3 region of the hippocampus, the neuron density in both infection and PBS groups declined with aging (Figure 3b&d). Compared to the controls, the infection group consistently showed lower neuron density at all five time points, and the difference between two groups increased from 4 Dpi to 450 Dpi. Specifically, the neuron density in the infection group was 7.0, 9.6, 17.0, 25.5, and 13.4 lower than the PBS group.

**Figure 3.**
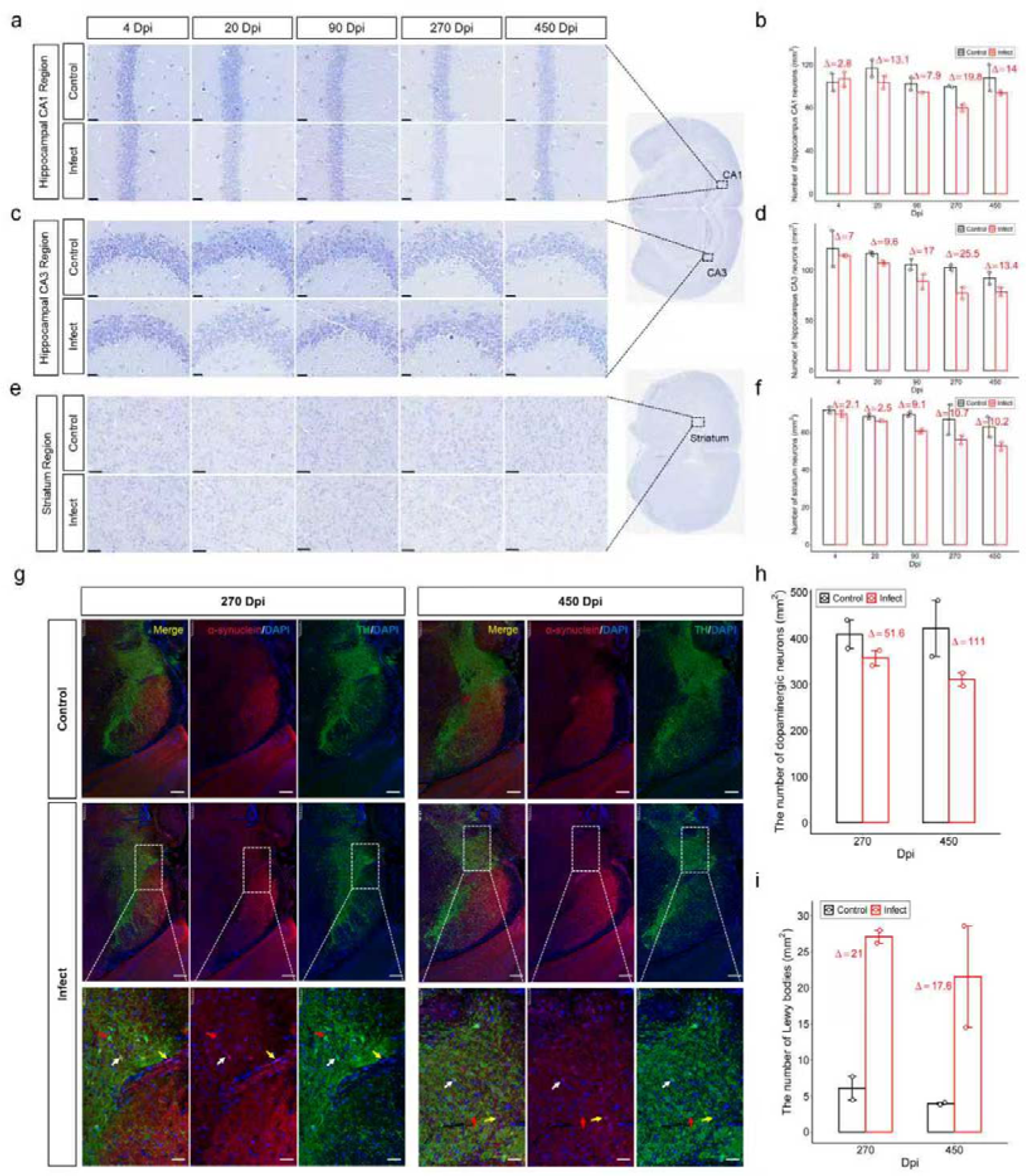
Neuronal loss and Parkinson’s disease-like neuropathological changes in the CNS of mice following influenza virus infection. (**a**), (**b**), and (**c**) refer to the images of the CA1 and CA3 regions of the mouse hippocampus (600×, Scale bar = 25 μm) and the striatum (400×, Scale bar = 50 μm), respectively, at five time points by Nissl staining. (**d**) and (**e**) refer to the density of neurons in the whole CA1 and CA3 regions of the mouse hippocampus, respectively, at five time points. (**f**) refers to the neuron density in the ROI of the mouse striatum at five time points. The bars show the average and the standard errors, and the delta indicates the average difference between the infection group and the PBS group. (**g**) refers to the images of the substantia nigra pars compacta (SNpc) at 270 and 450 Dpi, respectively, obtained by TH/α-synuclein double-labeling immunofluorescence (100×, Scale bar = 200 μm for low magnification; 400×, Scale bar = 50 μm for high magnification). (**h**) and (**i**) refer to the density of dopaminergic neurons and α-synuclein–positive aggregates, respectively, in the SNpc at 270 and 450 Dpi. n = 2 mice per group. Representative images are shown.

A similar phenomenon was observed in the striatum of infected mice, where the neuron density in the Region Of Interest (ROI) from the striatum was lower than that in the control group and the difference between two groups also increased from 4 Dpi to 450 Dpi (Figure 3e&f). Specifically, the neuron density in the infection group was 2.1, 2.5, 9.1, 10.7, and 10.2 lower than the control group. These results suggested significant reduction of neuron density in CNS of the middle and old-age mice after influenza virus infections.

### Influenza virus infection induces Parkinson’s-like neuropathological changes in aged mice

The above analysis indicates that influenza virus infection leads to neuronal loss in aged mice, which may impair cognitive function and contribute to the development of neurodegenerative disorders. Therefore, we further investigated whether influenza virus infection could trigger Parkinson’s disease (PD)-like pathology. Using TH/α-synuclein double-labeling immunofluorescence, we systematically assessed neurodegenerative changes in the substantia nigra (SN) of mice, focusing on two core pathological features of PD: the loss of dopaminergic neurons and the accumulation of α-synuclein–positive aggregates (Figure 3g). The results showed that at 270 Dpi (Figure 3h), the density of dopaminergic neurons in the substantia nigra pars compacta (SNpc) was significantly reduced (delta = 51.6 /mm^2^) compared to the PBS group. By 450 Dpi, the neuronal density in the PBS group remained stable, whereas the infected group exhibited a further decrease relative to both the control group (delta = 111 /mm^2^) and their own neuronal density at 270 Dpi (delta = 59.4 /mm^2^) (Figure 3h).

Lewy body-like cytoplasmic inclusions were observed in infected mice at both 270 and 450 Dpi (Figure 3g). Quantitative analysis revealed that the average density of α-synuclein–positive aggregates at 270 Dpi was 27/mm², compared to only 6/mm² in the PBS group—a three-fold increase (Figure 3i). By 450 Dpi, α-synuclein–positive aggregates density slightly decreased to 21/mm² but remained higher than that of the control group (4/mm²) (Figure 3i). These findings suggest that influenza virus infection may contribute to the development of Parkinson’s-like neuropathological changes in aged mice.

### Long-term transcription analysis in lung, brain and blood

To elucidate the molecular mechanisms underlying life-long multi-organ injury in mice following influenza virus infection, we employed RNA-Seq to profile the transcriptomes of the lungs, blood, and brain at five time points after infections. Differential gene expression analysis and functional enrichment analysis were subsequently performed (see Methods). As illustrated in Figure 4a, the lungs exhibited the highest number of differentially expressed genes (DEGs) (> 1,000), with DEGs detected across all five time points. However, these DEGs were mainly detected at 4, 20, and 90 Dpi. The blood displayed over 400 DEGs, primarily at 4, 270, and 450 Dpi, while the brain showed approximately 100 DEGs, with the majority appearing at 90 Dpi. Notably, DEGs displayed substantial heterogeneity across organs and time points, with most being uniquely associated with a specific organ and time point (Figure S1). Therefore, we next performed functional analysis of DEGs to delineate organ-specific responses to viral infections.

**Figure 4.**
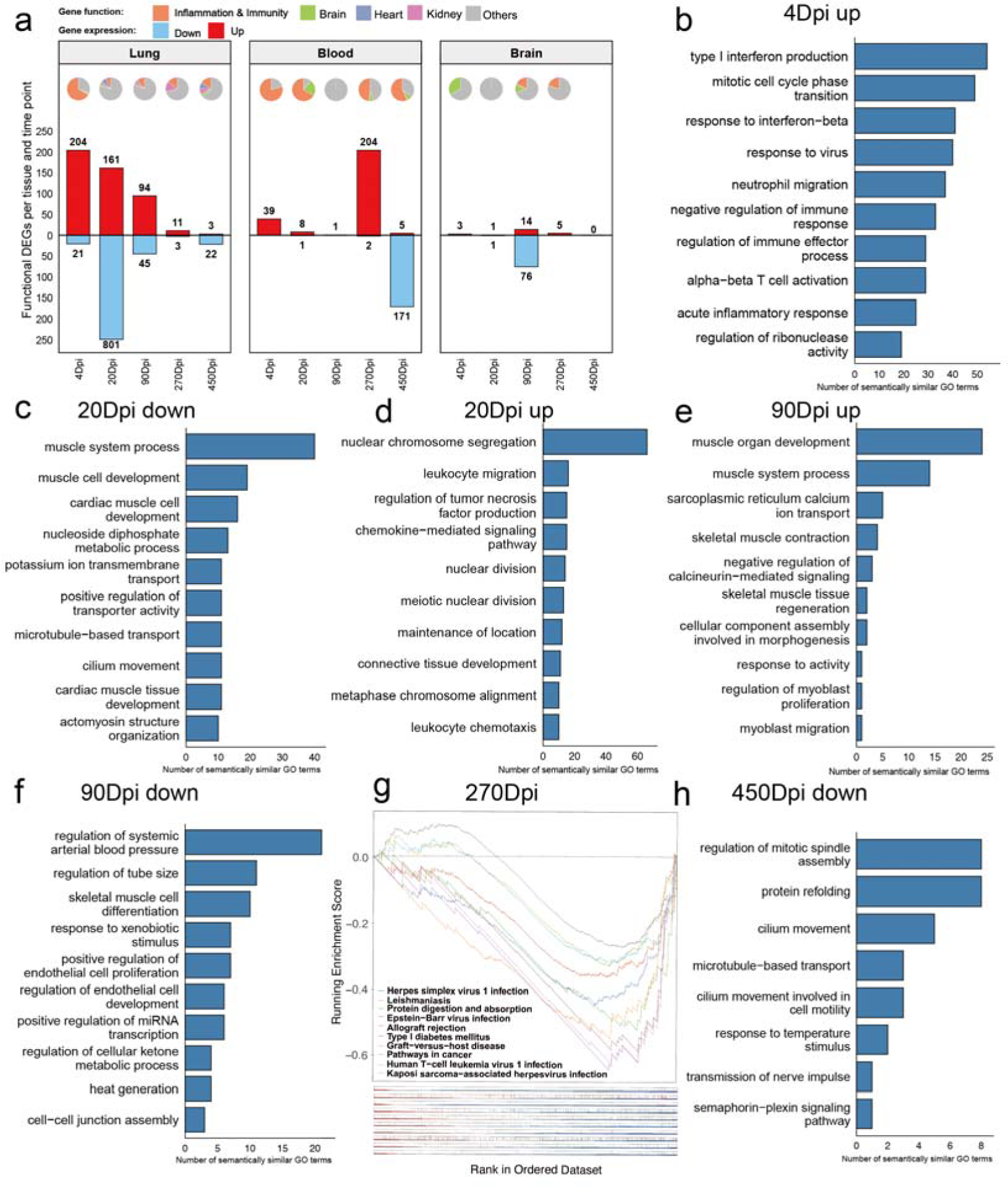
Differential gene expression analysis and functional enrichment analysis of the transcriptome in mice lung, blood and brain at five time points. (**a**) Number of DEGs identified in mice lung, blood and brain (bars) and functional composition of DEGs (pie charts) at five points. Subpanels of (**b**)-(**f**) and (**h**) refer to the enriched biological processes of up- and down-regulated DEGs in lung at different time points after infections. The GO terms were clustered, and each subpanel showed the number of GO terms in each cluster. (**g**) refers to the top 10 enriched KEGG pathways by GSEA analysis.

In the mice lung, 225 DEGs were identified at 4 Dpi, and the majority of them were upregulated (204 genes) (Figure 4a). Functional enrichment analysis revealed that upregulated genes were primarily associated with type I interferon (IFN-I) production, antiviral response, and inflammatory pathways, indicating a robust antiviral immune response (Figure 4b). No biological processes were enriched among downregulated genes. By 20 Dpi, 962 DEGs were detected, with a predominance of downregulated genes (801 genes) (Figure 4a). Downregulated genes were significantly enriched in pathways related to cilium movement, axoneme assembly, and ciliary function, suggesting potential impairment of pulmonary respiratory function (Figure 4c). In contrast, upregulated genes were linked to mitotic processes, including nuclear chromosome segregation and nuclear division, implying the initiation of tissue repair (Figure 4d). At 90 Dpi, the lungs still exhibited multiple DEGs (94 upregulated and 45 downregulated). Upregulated genes were predominantly involved in myocyte differentiation, muscle development, muscle contraction, and tissue morphogenesis, indicating activation of myogenic repair mechanisms (Figure 4e). Downregulated genes were enriched in processes such as blood pressure regulation, endocrine signaling, and thermoregulation, suggesting persistent metabolic and homeostatic disturbances (Figure 4f). By 270 Dpi, only a limited number of DEGs remained (11 upregulated and 3 downregulated). Gene Set Enrichment Analysis (GSEA) revealed upregulated pathways related to viral infection response such as Herpes simplex virus 1 infection and Epstein−Barr virus infection, implying residual immune activity (Figure 4g). At 450 Dpi, the lungs displayed 3 upregulated and 22 downregulated genes (Figure 4a). The upregulated genes including Ly6c, Ccl5, and Nkg7 were all immune-related. GSEA further confirmed enrichment of viral infection and immune-related pathways (Figure S2a), suggesting lingering inflammation. Downregulated genes were associated with mitosis, protein folding, and ciliary function, reinforcing the notion of sustained respiratory dysfunction (Figure 4h). These findings align with histopathological observations in aged mice, demonstrating persistent pulmonary impairment.

In the brain, only a limited number of DEGs were detected at 4 and 20 Dpi, primarily associated with functions of Neuroactive ligand-receptor interaction (Figure S2b&c). Notably, at 90 Dpi, a substantial number of DEGs emerged, consisting of 76 downregulated and 14 upregulated genes (Figure 4a). Downregulated genes were significantly enriched in brain developmental processes, including hindbrain development, midbrain development, and metencephalon development (Figure 5a), suggesting potential disruptions in neurogenesis. Additionally, synaptic transmission-related pathways such as chemical synaptic transmission and regulation of postsynaptic density assembly were suppressed. In contrast, the upregulated genes were predominantly involved in endocrine regulation, including regulation of gonadotropin secretion and steroid hormone secretion (Figure S2d), indicating dysregulation of the neuroendocrine axis at this stage. By 270 Dpi, only 5 DEGs (all upregulated) were observed in the brain (Figure 4a), including Drd2 which is implicated in dopamine binding. GSEA analysis identified 10 enriched pathways (Figure 5b), and all of them were upregulated. Three of them were linked to neurological functions including “Cocaine addiction”, “Dopaminergic synapse”, and “cAMP signaling pathway”. At 450 Dpi, no significant DEGs were detected in the brain (Figure 4a). However, GSEA revealed 6 enriched pathways (Figure 5c), four of which were associated with neuronal signaling and dysfunction, including “Neuroactive ligand-receptor interaction”, “Long-term depression” (downregulated), “Parkinson’s disease”, and “Synaptic vesicle cycle”. These findings suggest persistent neurofunctional perturbations long after viral clearance, potentially contributing to chronic neurological sequelae.

**Figure 5.**
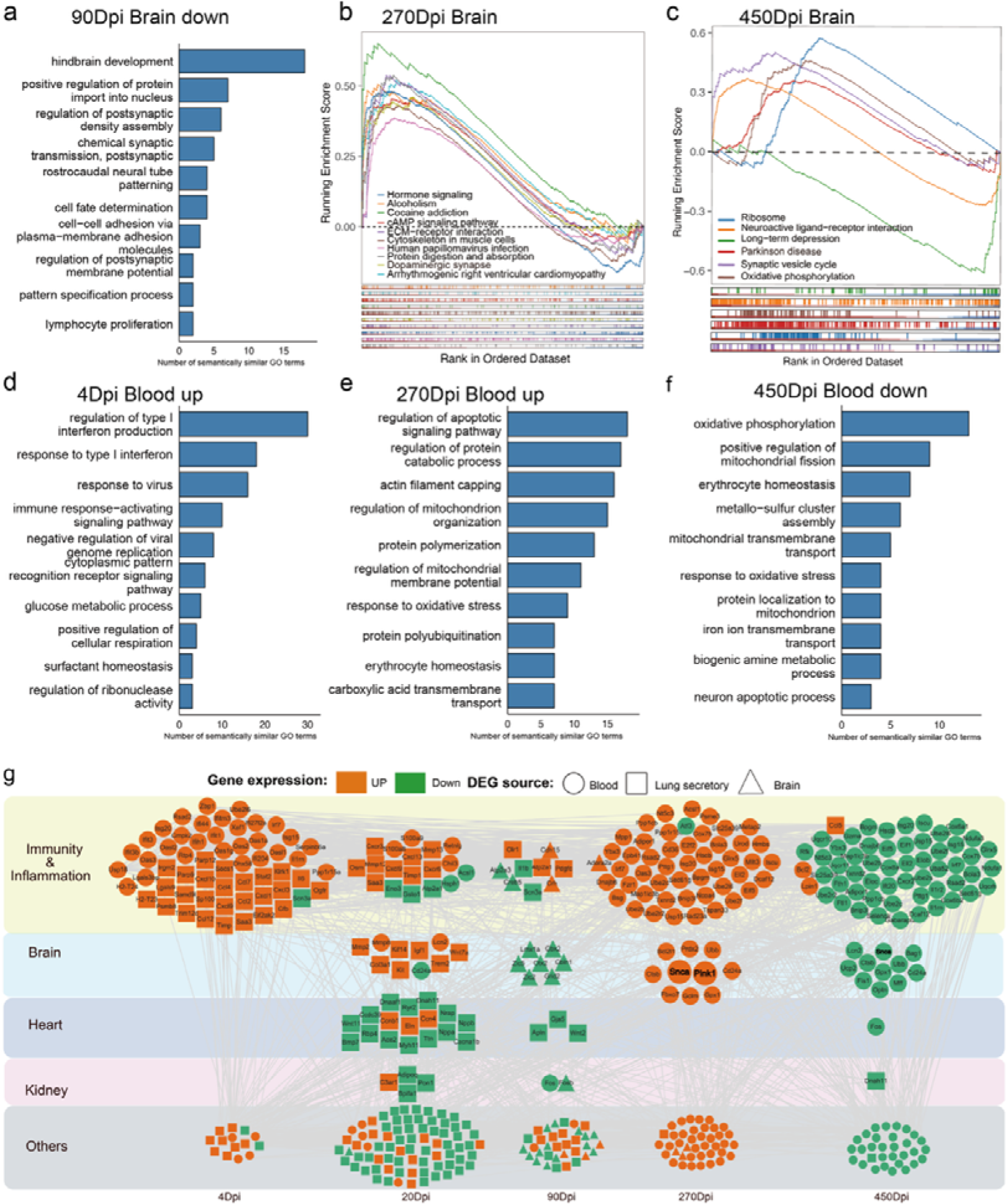
Transcriptome analysis and functional enrichment analysis in brain and blood. Subpanels **a**, **d**, **e** and **f** refer to enriched biological process of DEGs identified in brain or blood at given time points post infections. Subpanels **b** and **c** refer to enriched top 10 KEGG pathways of DEGs identified in brain at 270 and 450 Dpi by GSEA analysis. (**g**) Protein-protein interaction network between DEGs identified in the lung that encoded secreted proteins and the DEGs identified in brain and blood. The network was organized by time points post-infections. Nodes in orange and green refer to up-regulated and down-regulated DEGs, respectively. The shape of nodes refers to the source tissue of DEGs as indicated by the figure legend on the top. The nodes were manually clustered by functions.

In the blood, 39 DEGs were identified at 4 Dpi (Figure 4a), all of which were upregulated. Functional enrichment analysis revealed their involvement in antiviral response, regulation of viral genome replication, and type I interferon (IFN-I) signaling, mirroring the transcriptional profile observed in the lungs at this time point (Figure 5d). At 20 and 90 Dpi, only a limited number of DEGs were detected in the blood (Figure 4a), none of which showed significant enrichment for specific biological functions. Strikingly, by 270 Dpi, a substantial number of DEGs emerged (204 genes, predominantly upregulated) (Figure 4a). These genes were primarily associated apoptotic signaling, protein catabolism, mitochondrion function, oxidative stress responses, and erythrocyte homeostasis (Figure 5e), suggesting the presence of systemic chronic inflammation at this stage. By 450 Dpi, the blood exhibited 171 downregulated and 5 upregulated DEGs. The downregulated genes were significantly enriched in pathways related to oxidative phosphorylation and mitochondrial functions (Figure 5f), indicating dysregulation of cellular energy production and metabolic homeostasis. The few upregulated genes were linked to apoptosis and developmental processes in multiple tissues, including renal glomeruli, oocytes, and muscle fibers, potentially reflecting long-term multi-organ sequelae of the infection (Figure S2e).

### Systemic multi-organ effects mediated by pulmonary secreted proteins and inflammatory responses

Our pathological and transcriptomic analyses demonstrate that influenza virus infection induces long-term damage and transcriptional alterations not only in the lungs but also in multiple distal organs. We propose that pulmonary infection generates a cascade of effects that disseminate systemically through the bloodstream, ultimately impacting other organs. Functional characterization of the DEGs in mice lung identified multiple immune and inflammatory genes such as Isg15 and Cd24a that were persistently expressed across nearly all five time points (Figure 4a & Figure 5g). Notably, we also detected several DEGs related to cardiac and renal functions such as Nppa and Adipoq (Figure 4a and Figure 5g). DEGs identified in blood were predominantly enriched for immune and inflammatory factors such as Il6 and Il1, along with a subset of genes potentially influencing neurological functions such as Snca and Pink1 that were upregulated at 270 Dpi (Figure 4a and Figure 5g). Similarly, DEGs identified in the brain included both neural-specific genes such as Drd2 and immune mediators such as Crh observed at 90 Dpi (Figure 4a and Figure 5g). These findings collectively demonstrate the existence of a robust immune-inflammatory network interconnecting all three organs.

We hypothesized that the secretory proteins that were encoded by lung DEGs may be significantly altered post-infection. They can enter circulation and mediate distal organ effects. To investigate this, we constructed a protein-protein interaction network integrating 351 pulmonary secretory proteins encoded by lung DEGs with those encoded by DEGs identified in blood and brain tissues (Figure 5g). The network analysis revealed significant connectivity between pulmonary secretory proteins and those encoded by DEGs in both blood and brain tissues. Analysis of the spatiotemporal dynamics of the network showed distinct characteristics of the network at each time points. At early phase (4 Dpi), the protein-protein interaction network revealed a robust interplay between pulmonary and circulatory immune-inflammatory factors. Key mediators such as Cmpk2 and Dhx58 were highly upregulated in both lungs and blood, forming dense interaction clusters (Figure 5g). This pattern indicates acute systemic inflammation, likely driven by the initial viral load and host antiviral response. At the recovery phase (20 Dpi), organ-specific inflammation persisted. Viruses have been cleared from the mice at this stage (Figure 1d). Only four upregulated immune factors persisted in the blood, suggesting gradual resolution of systemic inflammation. However, the lungs retained multiple immune-related DEGs such as Cxcl9, consistent with persistent local inflammation observed histologically. Notably, several pulmonary secretory proteins at this stage were associated with disorders in distal organs: 17 genes were associated with cardiac disease and most of them were down-regulated such as Ttn and Nrap; 4 genes were associated with renal diseases and three of them were down-regulated such as Adipoq; 7 genes were associated with brain diseases and all of them were up-regulated such as Trem2 and Kif14. This suggests potential remote organ effects mediated by lung-derived factors even after viral clearance. During the intermediate phage (90 Dpi), neurological dysregulation emerges. While low-level immune activity persisted in the lungs such as Crh and Olr1, the brain exhibited widespread downregulation of neurofunctional genes such as Gbln1 and Zic5, alongside a subset of immune-modulatory genes such as Crh and Atp2a3, a renal disease related gene Fosb. Intriguingly, two immune genes, e.g., Atp2a3 and Atp2a1 that were identified in the brain and lung, respectively, interacted with each other, suggesting a potential lung-brain connection mediated by immunity. During the late phase (270 Dpi), a striking resurgence of immune-inflammatory genes occurred in the blood such as Irf7 and Isg15, accompanied by neurologically relevant genes such as Pink1 (mitophagy regulator) and Snca (encoding α-synuclein) (highlighted in Figure 5g). The upregulation of Snca, a key protein in Parkinson’s disease pathogenesis, suggests a plausible mechanism for post-viral neurodegenerative risk. During the chronic phase (450 Dpi), lots of immune genes such as Cox7b and Cd24a were downregulated in the blood, including those elevated at 270 Dpi, indicating immune exhaustion or resolution. Several neurofunctional genes such as Snca and Ctsb were also downregulated, aligning with neuron loss observed in aged-infected mice^14^.

## Discussion

Acute viral infections can cause severe illness in hosts^15^. Current research primarily focuses on the acute and recovery phases of viral infection, with limited investigation into the sequelae of viral infections. Long COVID has raised awareness that viral infections may have long-term effects on hosts^16,17^. This study, to the best of our knowledge, is the first to investigate the lifelong impacts of influenza virus infection in mice, revealing that infection in juvenile mice can lead to lifelong pathological damage in the lungs, heart, and kidneys, as well as neuronal loss and even Parkinson’s disease-like pathological changes in old age. These findings suggest that severe influenza virus infections have more profound and lasting effects on hosts than previously anticipated, underscoring the importance of timely vaccination or early intervention after infection to reduce the occurrence of severe viral infections.

Existing studies have mainly focused on the acute phase and recovery period, with the longest follow-up extending to six months post-infection^18,19^. For example, Hosseini et al. reported that infection with neurotropic (H7N7) and non-neurotropic (H3N2) influenza A virus strains induced impairments in learning, memory, and behavior in mice, which were still detectable at 120 days post-infection^20^. These changes were associated with long-term hippocampal neuroinflammation, synaptic loss, and reduced synaptic plasticity. Our study is the first to demonstrate that influenza virus infection can cause lifelong effects. In contrast to previous work, we did not observe significant transcriptomic alterations in mice brain during the acute or early phases of infection, but we detected pronounced effects at three months post-infection and beyond. This discrepancy may arise because our transcriptomic analysis encompassed the whole brain rather than focusing on specific regions like the hippocampus. Additionally, we observed significant damage in other organs such as the heart and the damage persisted long after infection and increased with age^21^. One possible explanation is that severe viral infections in early life cause permanent damage to certain cells or tissues^22^. For instance, persistent pulmonary fibrosis may impair respiratory function, and such damage may accumulate over time, increasing the host’s susceptibility to disease^23^. Previous works which analyzed electronic medical records, showed that early-life viral infections, including influenza virus and herpesviruses, may increase the risk of neurodegenerative diseases such as Alzheimer’s disease (AD) and Parkinson’s disease (PD) in old age, consistent with our findings and supporting the notion that viral infections can have lifelong consequences for hosts^6,24^.

Investigating the long-term effects of viral infections on hosts is challenging due to numerous confounding factors^25^. Throughout their lives, humans experience various injuries, including infections (viral, bacterial, fungal, etc.), physical and chemical damage, and psychological stress, making it difficult to isolate the long-term impact of a specific viral infection^26^. Current research typically relies on epidemiological surveys, using large cohort studies to establish control groups and minimize confounding effects, as seen in the aforementioned work by Bjornevik et al^27^. However, this approach still struggles to exclude the influence of other factors. Our study leverages the advantages of mouse models: their relatively short lifespan allows for the investigation of lifelong effects; their housing in specific pathogen-free (SPF) environments greatly reduces environmental variables; and strict controls can be implemented to minimize confounding factors. Although low levels of viral and bacterial pathogens were detected in the lungs and blood of mice via transcriptomic analysis, these were likely sequencing contaminants. Moreover, since stringent controls were set in each time point, the impact of confounding factors on the mice was negligible. Thus, the lifelong effects of viral infection observed in our study are highly reliable.

Viral infections can affect multiple systems throughout the body, as exemplified by SARS-CoV-2, which causes damage to various organs^28^. Our study similarly revealed systemic effects, with severe damage in the lungs and varying degrees of long-term injury in the heart, kidneys, and brain. A recent study by Zheng et al. demonstrated that influenza virus can spread via the bloodstream to multiple organs, directly causing damage^29^. In contrast, our RNA-Seq data detected influenza virus sequences only in the lungs at 4 Dpi and no influenza viruses was detected in any other organs or at any other time points, suggesting that the observed multi-organ damage may not result from direct viral infections. This difference may be caused by strain heterogeneity. Based on transcriptomic analysis, we hypothesize that influenza virus infection alters the lung expression profile, leading to the release of secretory proteins into the bloodstream that directly or indirectly (via immune and inflammatory factors) damage other tissues and organs. For example, we observed significant inflammatory injury in the heart and kidneys, as well as differentially expressed inflammatory factors in the brain. Our analysis showed the existence of a robust immune-inflammatory network interconnecting all three organs. The network is characterized by i) core immune/inflammatory factors originating from the infected lungs; ii) systemic propagation through circulatory factors; iii) secondary immune activation in distal organs; iv) potential cross-talk between immune and organ-specific functional pathways. This interconnected network may explain the long-term multi-organ sequelae observed following pulmonary influenza infection, suggesting that pulmonary-derived factors sustain systemic inflammation that subsequently drives chronic dysfunction in distal organs.

Parkinson’s disease (PD) is a progressive neurodegenerative disorder characterized by the degeneration of dopaminergic neurons in the substantia nigra and the presence of Lewy bodies^30^. The pathogenesis of PD is diverse and complex, one contributing factor being viral infections. Previous studies have identified associations between various viral infections including influenza virus, HSV-1 and VZV, and Parkinson’s disease, but these were based solely on epidemiological investigations^31–33^. For example, Shirin Hosseini have reported higher incidences of PD in populations previously infected with influenza viruses^33^. This study is the first to demonstrate experimentally in mice that influenza virus infection during early life leads to two core neuropathological features of Parkinson’s disease in old age: loss of dopaminergic neurons in the substantia nigra pars compacta and accumulation of Lewy body–like α-synuclein inclusions. Additionally, significant neuronal reduction was observed in the hippocampus and striatum. These findings indicate that mice infected with influenza virus during early life develop severe neurological impairment and Parkinson’s-like pathology in old age. Transcriptomic analysis suggests this damage may initiate as early as 90 Dpi, evidenced by substantial downregulation of brain development-related genes. At 270 days post-infection, multiple Parkinson’s-associated genes were upregulated in blood, particularly the α-synuclein gene. Correlated with brain pathology, these changes suggest Parkinson’s-like pathological alterations were already developing at this stage. This implies influenza virus infection triggers Parkinson’s disease pathogenesis much earlier than previously assumed.

Several limitations should be noted. First, while we observed Parkinson’s-like pathology and neuronal loss, the lack of behavioral tests prevents definitive confirmation of Parkinson’s disease diagnosis or assessment of cognitive and motor impairments. Second, given the long timeline from childhood to old age, our study only examined five timepoints, leaving gaps in understanding interim changes. Future studies should incorporate more timepoints. Third, this work focused on select organs (lungs, brain, blood), leaving other critical organs (liver, spleen, gut) unexamined regarding viral infection effects. Fourth, while transcriptomics revealed molecular changes in lungs, brain and blood, the detailed molecular mechanisms underlying multi-organ effects and long-term health impacts remain unclear. Comprehensive multi-omics approaches (proteomics, metabolomics, epigenomics, metagenomics) are needed to fully elucidate these mechanisms.

This study represents a significant exploration into the long-term effects of influenza virus infection in mice. While we have obtained meaningful and intriguing findings, our work also opens new avenues for investigating the prolonged consequences of viral infections, with numerous questions warranting further research. First, our study focused on the impact of influenza infection in juvenile mice. It remains unknown how infection at other life stages—such as adolescence, adulthood, or even old age—might influence disease susceptibility and overall health. Second, our findings demonstrate that influenza infection leads to severe long-term consequences in mice. An important follow-up question is whether preventive measures, such as influenza vaccination, or prompt antiviral treatment post-infection could mitigate these effects. Third, beyond influenza virus, do other viral infections—such as SARS-CoV-2 or respiratory syncytial virus (RSV)—also induce similarly severe outcomes, including Parkinson’s disease-like pathology? Fourth, the viral dose used in this study was relatively high, resulting in severe infection. Would a lower viral dose produce milder or negligible long-term effects? These unresolved questions highlight the need for expanded research to better understand the lifelong health implications of viral infections and develop effective intervention strategies.

## Conclusion

This study is the first to demonstrate that influenza virus infection can cause lifelong damage to multiple organs in mice and induce Parkinson’s disease-like pathological changes in their later life. Transcriptomic analysis revealed that influenza virus infection in the lungs may exert systemic effects through secreted proteins and inflammatory responses. The findings highlight the lifelong consequences of viral infections on the host, underscoring the need to mitigate severe viral infections, which would have a far-reaching impact on the prevention and treatment of viral infectious diseases.

## Methods

### Animal model establishment and viral infection

Six-week-old female C57BL/6J mice (18–22 g) were purchased from Hunan SJA Laboratory Animal Co., Ltd (Changsha, China, license number: SCXK(Xiang)2021-0002). Mice were housed in a specific pathogen-free (SPF) facility under standardized environmental conditions (22 ± 1°C, 50 ± 10% relative humidity, 12-hour light/dark cycle) and acclimatized for 1 week. A total of 50 mice were randomly assigned based on body weight to either the infection group (n = 25, 5 mice per time point) or the control group (n = 25, 5 mice per time point)^34^.

The mouse-adapted influenza A virus strain A/Hong-Kong/8/1968 (H3N2) was propagated in 10-day-old embryonated chicken eggs and titrated to a final concentration of 3.53×10 TCID^35^.

### Virus infection and sample collection

Mice were anesthetized by intraperitoneal injection of sodium pentobarbital (50 mg/kg). In the infected group, 50 μL of viral suspension (0.5 LD_50_) was administered intranasally. Control mice received 50 μL of sterile phosphate-buffered saline (PBS) containing 0.1% bovine serum albumin (BSA). Body weight was monitored daily, and mice exhibiting >30% weight loss were euthanized. At each designated time point, five mice were sacrificed for tissue collection. Mice were euthanized under deep anesthesia, and tissues were rapidly harvested without transcranial perfusion. For brain tissue, three mice were selected for RNA-Seq analysis: their brains were snap-frozen in liquid nitrogen for 5 minutes and stored at −80°C. The remaining two mice had their brains perfused with PBS and fixed in 4% paraformaldehyde (PFA) for Nissl and immunofluorescence staining. For lung tissue, four mice were selected. Their left lungs were snap-frozen in liquid nitrogen for 5 minutes and stored at −80°C for RNA-Seq, while the right lungs were washed with PBS and fixed in 4% PFA for hematoxylin and eosin (H&E) staining and Masson’s trichrome staining. The hearts and left kidneys from the same four mice were washed with PBS and fixed in 4% PFA for histological staining. Blood was collected via cardiac puncture (approximately 1.5 mL per mouse) and mixed with TRIzol™ reagent at a 1:3 volume ratio before storage at −80°C for RNA-Seq.

### Lung viral load quantification at 4 days post-infection

To assess viral replication, lung tissues from infected mice at 4 Dpi were homogenized in 1 mL of ice-cold PBS using a sterile tissue grinder. The homogenates were centrifuged at 4°C, 12,000 × g for 10 minutes, and the supernatants were subsequently collected. Madin-Darby Canine Kidney (MDCK) cells were then seeded into 96-well plates and incubated with ten-fold serial dilutions of lung homogenates. After incubation for 72 hours, cytopathic effects (CPE) were evaluated under the microscope, and the 50% tissue culture infectious dose (TCID) was calculated using the Reed–Muench method^34^.

### Hematoxylin and Eosin (H&E) Staining and Masson’s trichrome staining

Lung, heart, and kidney tissues were collected from C57BL/6 mice and immediately fixed in 4% PFA at 4℃ for 24 hours. The fixed tissues were dehydrated in a graded ethanol series (70%, 80%, 90%, 95%, and 100%, 1 hour each per step), cleared in xylene (twice, 15 minutes each), and embedded in paraffin. Paraffin-embedded tissues blocks were sectioned at a thickness of 5 μm using a rotary microtome (Leica RM2235, Leica Biosystems, Germany). Tissue sections were deparaffinized in xylene (twice, 10 minutes each), rehydrated through a descending ethanol series (100%, 95%, 90%, 80%, and 70%, 5 minutes each), and rinsed in distilled water.

For hematoxylin and eosin (H&E) staining, sections were immersed in hematoxylin for 5 minutes, differentiated in 1% acid ethanol for 30 seconds, rinsed under running tap water for 5 minutes, and then counterstained with eosin for 2 minutes^36^.

For Masson’s trichrome staining, sections were stained with Weigert’s iron hematoxylin for 10 minutes, rinsed in tap water for 10 minutes, then stained with Biebrich scarlet-acid fuchsin solution for 10 minutes. After differentiation in phosphomolybdic-phosphotungstic acid solution for 10 minutes, sections were stained with aniline blue for 5 minutes and treated with 1% acetic acid for 2 minutes^37^.

Sections were then dehydrated, cleared, and mounted with neutral balsam. Histological images were captured using a slide scanning system (Pannoramic SCAN II, Hungary).

### Lung injury assessment

Microscopic images were acquired using an Olympus BX53 microscope equipped with a DP74 digital camera. Uniform settings for brightness, contrast, and threshold were applied during image acquisition and analysis to ensure consistency across samples. Lung injury was semi-quantitatively evaluated on H&E-stained lung tissue sections by two blinded pathologists based on six histological features: alveolar edema, cell debris, inflammatory cell infiltration, alveolar epithelial cell infiltration, hemorrhage, and alveolar wall thickening. Each parameter was scored on a scale from 0 to 4, where 0 indicated no abnormality, 1 indicated minimal changes, 2 indicated mild changes, 3 indicated moderate changes, and 4 indicated severe or extensive damage. This results in a total injury score ranging from 0 to 24. For each H&E-stained section, five regions of interest (ROIs) were selected from representative regions, including the alveolar area, perivascular space, and peribronchiolar regions, to ensure a comprehensive assessment of lung pathology.

To measure the pulmonary fibrosis in mice, five pathological ROIs were randomly selected on each section. ROIs were specifically chosen from alveolar regions distant from large airways and blood vessels to avoid interference from naturally collagen-rich structures. The collagen content was calculated based on Masson-stained lung sections by quantifying the percentage of collagen-positive area within alveolar regions with ImageJ software (NIH, USA), using standardized color thresholding parameters^38^.

### Cardiac injury assessment

Cardiac injury was assessed semi-quantitatively on H&E-stained heart tissue sections based on three key histopathological parameters: inflammatory cell infiltration, myocardial fiber degeneration, and myocardial fibrosis. For each parameter, severity was scored on a scale from 0 to 4. The total cardiac injury score ranged from 0 to 12. Histological evaluation was performed independently by two blinded pathologists to ensure objectivity and consistency. Sections were examined in five non-overlapping high-power fields (HPFs) selected from representative regions of the left ventricular wall, interventricular septum, and subendocardial area to ensure comprehensive assessment of myocardial injury.

### Renal injury assessment

Renal damage was evaluated on H&E-stained kidney sections based on two histopathological features: inflammatory cell infiltration in glomeruli and glomerulosclerosis. A minimum of 30–50 glomeruli per animal were examined. For each animal, the number of glomeruli showing inflammatory cell infiltration and the number of sclerotic glomeruli were recorded. The percentage of affected glomeruli was calculated separately for each parameter, and the mean percentages were used to indicate the severity of renal injury. All assessments were conducted by two blinded pathologists.

### Nissl staining and neuronal quantification

Brain tissues were collected without transcardial perfusion and fixed by immersion in 4% paraformaldehyde (PFA), dehydrated, embedded in paraffin, and coronally sectioned at a thickness of 5 μm using a rotary microtome (Leica RM2235, Leica Biosystems). Every sixth section (sampling interval = 120 μm) encompassing the hippocampus and striatum was selected for analysis, and three anatomically matched sections were assessed per animal. For Nissl staining, sections were deparaffinized, rehydrated, and stained with 0.1% cresyl violet solution at room temperature for 15 minutes. Differentiation was performed in 95% ethanol containing 0.1% glacial acetic acid, followed by dehydration in graded ethanol and clearing in xylene. Sections were then cover slipped with neutral resin^39^.

Neuronal quantification was performed manually in the whole CA1 and CA3 subfields of the hippocampus and the ROIs of dorsolateral region of the striatum. ROIs were delineated bilaterally based on a standard mouse brain atlas. Only Nissl-positive neurons with clear, round nuclei and prominent nucleoli were counted. Cell counting was performed by two blinded observers to reduce bias. Image analysis was conducted using ImageJ software (NIH), with consistent thresholding parameters applied across all samples. Neuronal density was expressed as the number of neurons per mm².

### Immunofluorescence staining and quantification of TH□ neurons and **α**-Synuclein–Positive Aggregates

The tissue sections were deparaffinized, rehydrated, and rinsed in buffer to antigen retrieval. Antigen retrieval was carried out in 10 mM sodium citrate buffer (pH 6.0) at 121 °C for 15 minutes under high pressure. After cooling, sections were blocked with 5% normal donkey serum and incubated overnight at 4 °C with rabbit anti-tyrosine hydroxylase (TH) antibody (Abcam, ab112, 1:500) and mouse anti-α-synuclein antibody (CST #4179, 1:1000). After PBS washing, sections were incubated at room temperature for 1 hour with Alexa Fluor 488-conjugated donkey anti-rabbit (Invitrogen, A-11008, 1:1000) and Alexa Fluor 594-conjugated donkey anti-mouse (Invitrogen, A-11005, 1:1000) secondary antibodies. Nuclei were counterstained with DAPI (1 μg/mL, Sigma) for 5 minutes, and sections were mounted using ProLong™ Gold antifade reagent.

Confocal images were acquired using a Zeiss LSM 880 microscope. Z-stack images were collected at 0.5 μm intervals using ZEN software (version 3.3). For each animal, three anatomically matched coronal brain sections containing the substantia nigra pars compacta (SNpc) were analyzed.

TH neurons were manually counted throughout the entire SNpc using 3D reconstructions generated from the Z-stack image sets. Neurons were identified based on cytoplasmic TH immunoreactivity and the presence of a visible nucleus. α-Synuclein–Positive Aggregates were defined as large, intracytoplasmic α-synuclein-positive inclusions located within TH neurons. α-Synuclein–Positive Aggregates were manually identified and quantified throughout the entire SNpc. The total number of TH neurons a was recorded. The density of α-synuclein aggregates was expressed as the number of particles per mm²^40^.

### RNA extraction, library preparation, and RNA-Seq

Total RNA was extracted from approximately 3 μL of homogenized lung, brain, and blood tissues using TRIzol reagent (Invitrogen), followed by DNase I (Takara) treatment to remove genomic DNA contamination. RNA quality was assessed using a NanoDrop spectrophotometer (A260/A280 ≥ 1.8) and Agilent 2100 Bioanalyzer (RNA Integrity Number, RIN ≥ 7.0).

Polyadenylated [poly(A)+] RNA was enriched using Oligo(dT) magnetic beads and fragmented. First-strand and second-strand cDNA synthesis was performed using SuperScript IV reverse transcriptase (Invitrogen). Sequencing libraries were constructed using the NEBNext Ultra II DNA Library Prep Kit (New England Biolabs) following the manufacturer’s instructions. Paired-end sequencing (2 × 150 bp) was performed on the Illumina NovaSeq 6000 platform, generating approximately 30 million reads per sample.

### RNA-Seq data processing and differential gene expression analysis

Raw sequencing reads were trimmed for adapters and low-quality bases using fastp (v0.23.2)^41^. Clean reads were aligned to the Mus musculus reference genome (GRCm39) from NCBI using HISAT2 (v2.2.1) with default parameters^42^. Gene-level read counts were obtained using FeatureCounts (v2.0.1)^43^. To evaluate global transcriptomic variation across samples and to identify potential outliers or batch effects, principal component analysis (PCA) was performed using the prcomp function in R (v4.3.1). Raw count data were first transformed using variance-stabilizing transformation (VST) as implemented in the DESeq2 package (v1.44.0). The top principal components were used to visualize clustering patterns by tissue type and infection status. Outliers were removed for further analysis (Figure S3). Differentially expressed genes (DEGs) were identified using DESeq2(v1.44.0) in R. Genes with an adjusted p-value < 0.05 and absolute log (fold change) > 1 were considered statistically significant^44^.

### GO enrichment analysis and GSEA

Gene Ontology (GO) enrichment analysis was conducted using the clusterProfiler R package (v4.12.6)^45^. GO terms with p < 0.05 and q < 0.05 were considered significantly enriched. For clarity, GO terms were clustered based on their semantic similarity using rrvgo (v1.16.0)^46^. Gene Set Enrichment Analysis (GSEA) was conducted using the gseKEGG function^45^ based on log (fold change)–ranked gene lists. Enrichment results were visualized with GseaVis (v0.1.1)^47^.

### Protein–protein interaction (PPI) network construction

The DEGs identified in mice lungs that encoded secreted proteins (curated from the Secreted Protein Database)^48^ and those identified in blood and brains were used in building the protein-protein interaction network using the STRING database (v11.5)^49^ with a confidence score threshold of > 0.7. The resulting network was visualized using Cytoscape software (v3.10.3)^50^. DEGs involved in BPs related to heart, brain, and kidney functions were considered to be associated these functions. The immune-inflammation genes were curated from two predefined gene sets (HALLMARK_INFLAMMATORY_RESPONSE and M7) in the Molecular Signatures Database (MSigDB)^51^.

### Virus and bacteria identification from RNA-Seq data

To identify potential virus and bacterial pathogens from mouse transcriptome data, first, host reads were removed by HISAT2 (v2.2.1)^52^; then, ribosomal RNA sequences were removed by aligning non-host reads to the SILVA database (release 138.2)^53^ using BLASTN (v2.16.0)^54^ with the e-value threshold of 10. The remaining reads were then queried the NT database ^55^using BLASTN with the e-value threshold of 10, and only the best-hits was retained for each read. To ensure reliable identification, several filtering criteria were applied: (i) hits with < 80% alignment coverage or < 90% sequence identity were discarded; (ii) reads assigned to non-viral and non-bacterial taxa were excluded; (iii) hits from endogenous viruses, plant viruses or bacteriophages were excluded. Taxonomic assignments were determined based on the taxonomy of the best hit. The abundance of virus or bacteria was quantified by RPKB (Reads Per Kilobase per Billion host reads)^56^, which was calculated as:

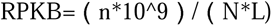

where n refers to the number of RNA-Seq reads assigned to a certain virus or bacterium, N refers to the number of reads mapped to the mouse genome (obtained from HISAT2 alignment), and L refers to the length of the genome of each corresponding virus species (retrieved from NCBl).

### Statistical Analysis

All statistical analyses were performed using GraphPad Prism 9.0 (GraphPad Software) and R software (v4.3.2). Each data point represents an independent biological replicate from a distinct animal. Sample sizes (n) for each group are indicated in the figure legends. Data are presented as mean ± s.e.m. unless otherwise stated.

Comparisons between two groups were performed using unpaired two-tailed Student’s t-tests were used^56^. Data distribution was assumed to be normal, but this was not formally tested due to the small sample sizes. No covariates were included in the statistical analyses, as all mice were age- and sex-matched and housed under identical experimental conditions. Exact P values are reported in the figures. Test statistics (t values), degrees of freedom, and sample sizes are provided where appropriate. Effect sizes (e.g., Cohen’s d) were not calculated in this study due to the small sample sizes. No Bayesian analyses or hierarchical/repeated-measures designs were performed.

## Ethics statement

All animal procedures were reviewed and approved by the Animal Ethics Committee of Hunan University (Approval No. HNUBIO202402001) and conducted in strict accordance with the ARRIVE guidelines, the NIH Guide for the Care and Use of Laboratory Animals, and the Institutional Animal Care and Use Committee (IACUC) guidelines.

## Data availability

The raw sequence data reported in this paper have been deposited in the Genome Sequence Archive, China National Center for Bioinformation / Beijing Institute of Genomics, Chinese Academy of Sciences (GSA: CRA029364) that are publicly accessible at at https://ngdc.cncb.ac.cn/gsa.

## Funding

This study was supported by the R&D Program of Guangzhou National Laboratory (GZNL2024A01002), National Natural Science Foundation of China (32170651 & 32370700), Hunan Provincial Natural Science Foundation of China (2024JJ2015), and the National Key Research and Development Program of China (2022YFC2303800).

## Acknowledgements

We are grateful to the members of Peng Lab for their insightful discussions on the manuscript and to Deng Lab for their valuable assistance and support throughout this study. We are also grateful to Dr. Lingxiao Zhang (Department of General Surgery, Xinhua Hospital, Shanghai Jiao Tong University School of Medicine) and Dr. Youwei Guo (Department of Neurosurgery, Xiangya Hospital, Central South University) for their insightful comments and critical revision of this manuscript.

## Competing interests

The authors declare that they have no competing interests.

## Authors’ contributions

Study design and supervision: YP and LD; Animal experiments and viral infection experiments: CL and LD; Pathological experiment and image analysis: RY, QH and GD; Transcriptomic data analysis: RY, XW and ZX; Manuscript draft: YP and RY; Manuscript review and revision: RY, YP, LD, ZM, TJ and DW; Funding: YP and ZM.

## Notes

### Competing Interest Statement

The authors have declared no competing interest.

