## Supplementary material for "Flu Virus Infection in Juvenile Mice Leads to Lifelong Multi-Organ Damage and Parkinsonian Pathological Changes in Aging": Figure S1

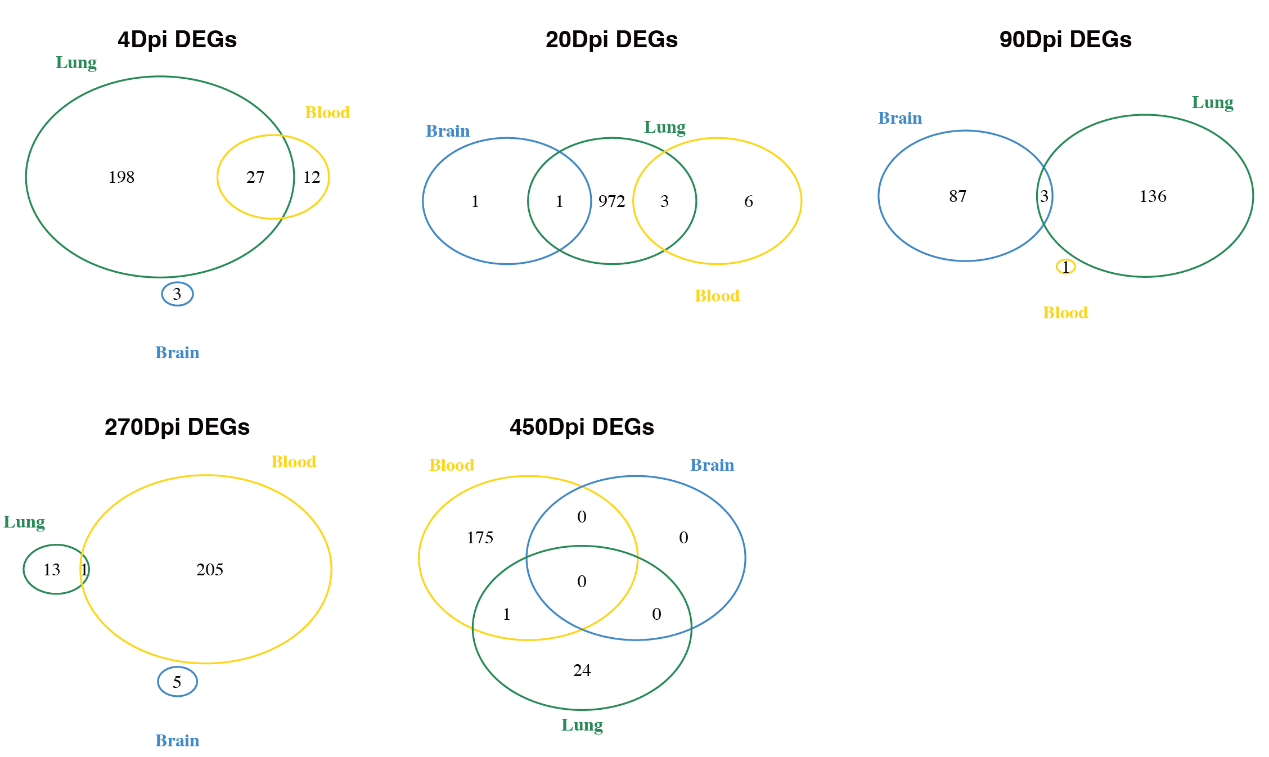

**Figure S1. Venn diagrams of differentially expressed genes (DEGs) in lung, blood, and brain at different time points post-infection.** Venn diagrams show the overlap of DEGs among lung (green), blood (yellow), and brain (blue) at five time points: 4 (Dpi), 20 Dpi, 90 Dpi, 270 Dpi, and 450 Dpi. Each number represents the count of DEGs specific to a tissue or shared between tissues at the indicated time point.
