## Supplementary material for "Flu Virus Infection in Juvenile Mice Leads to Lifelong Multi-Organ Damage and Parkinsonian Pathological Changes in Aging": Figure S2

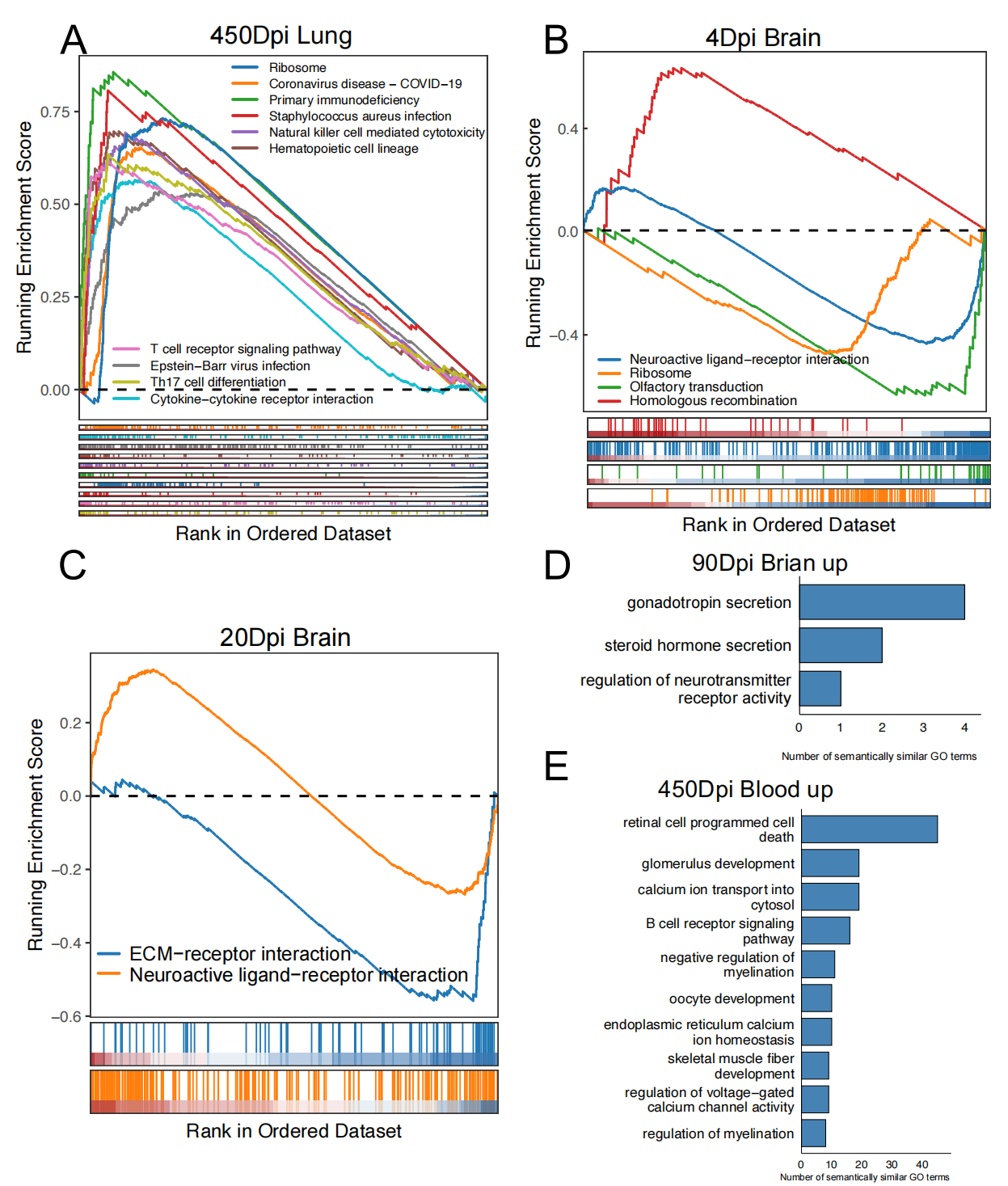

**Figure S2. Functional enrichment analysis of DEGs in lung, brain, and blood at different time points post-infection.** (a) Gene set enrichment analysis (GSEA) of lung DEGs at 450 Dpi showing enrichment in immune-related and infection-related pathways, including T cell receptor signaling, NK cell cytotoxicity, and hematopoietic cell lineage.(b) GSEA of brain DEGs at 4 Dpi, highlighting enrichment in pathways such as neuroactive ligand–receptor interaction, ribosome, olfactory transduction, and homologous recombination. (c) GSEA of brain DEGs at 20 Dpi, indicating enrichment of ECM–receptor interaction and neuroactive ligand–receptor interaction pathways. (d) Gene Ontology (GO) enrichment of upregulated genes in the brain at 90 Dpi, with terms related to hormone secretion and neurotransmitter receptor regulation. (e) GO enrichment of upregulated genes in blood at 450 Dpi, with terms associated with immune regulation, calcium signaling, programmed cell death, and development.
