## Supplementary material for "Flu Virus Infection in Juvenile Mice Leads to Lifelong Multi-Organ Damage and Parkinsonian Pathological Changes in Aging": Figure S3

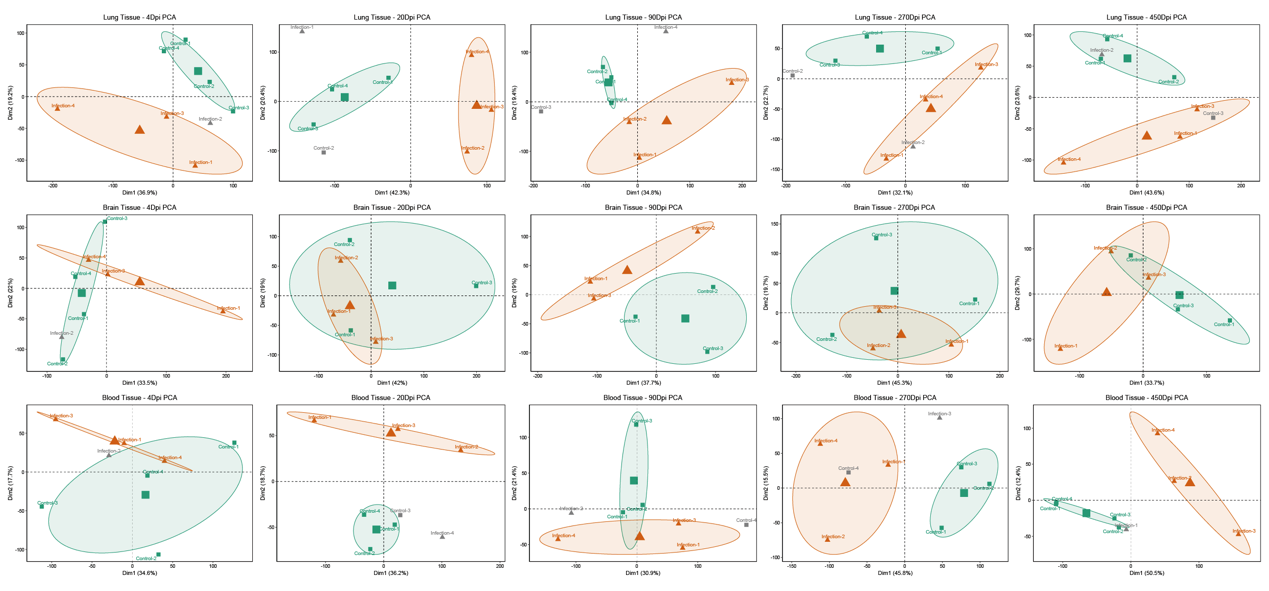

**Figure S3. Principal component analysis (PCA) of transcriptome profiles from lung, brain, and blood at different time points post-infection.** PCA plots of RNA-seq data from lung (top row), brain (middle row), and blood (bottom row) tissues at 4, 20, 90, 270, and 450 Dpi. Each point represents an individual sample, with infection groups (orange triangles) and control groups (green squares) shown separately. Gray points indicate samples that were removed from subsequent analyses. Ellipses indicate 95% confidence intervals for each group. Clear separations between infection and control samples at multiple time points indicate distinct transcriptional responses across tissues during both acute and long-term infection phases.
